# Perturbation of ubiquitin homeostasis promotes macrophage oxidative defenses

**DOI:** 10.1101/276964

**Authors:** Marie-Eve Charbonneau, Karla D. Passalacqua, Susan E. Hagen, Hollis D. Showalter, Christiane E. Wobus, Mary X.D. O’Riordan

## Abstract

Innate immune responses rely on specific pattern recognition receptors that induce downstream signaling cascades and promote inflammatory responses. Emerging evidence suggests that cells may also recognize alterations in cellular processes induced by infection. Protein ubiquitination is a post-translational modification essential for maintaining cellular homeostasis, and infection can cause global alterations in the host ubiquitin proteome. Here we used a chemical biology approach to perturb the cellular ubiquitin proteome as a simplified model to study the direct effect of ubiquitin homeostasis on macrophage responses. We show that perturbation of ubiquitin homeostasis results in a rapid and transient burst of reactive oxygen species (ROS) that promotes macrophage anti-infective capacity. ROS production was dependent on the activity of the phagocyte NADPH oxidase NOX2 and was associated with an increase in intracellular calcium. Our findings suggest that major changes in the host ubiquitin landscape may be a potent signal to rapidly deploy innate immune defenses.

---

The innate immune system provides rapid protective responses to various stimuli including injury and infection. To promote an effective inflammatory response, immune cells can initiate signaling cascades through pattern recognition receptors (PRR), which recognize diverse exogenous pathogen-associated molecular patterns (PAMP) or endogenous damage-associated molecular patterns (DAMP) (1). Although PAMP sensing can mediate protection from infection, many pathogens are able to avoid or suppress recognition by specific PRR, suggesting an advantage for the host to recognize other signals associated with infection (2). Moreover, the classic PRR model does not explain immune discrimination between pathogenic and non-pathogenic microbes. Indeed, emerging data suggest that the innate immune system can detect conserved pathogen-induced processes such as changes in the activation state of small Rho GTPases caused by microbial infection (2–4). More recently, the term HAMP (homeostasis-altering molecular processes) has been proposed to reflect that cells detect changes in cytoplasmic homeostasis induced during infection (5), such as activation of the inflammasome sensor NLRP3, triggered in response to a wide range of signals with no structural homologies. The HAMP-sensing hypothesis is analogous to the “guard hypothesis” in plant immunity, where disruption of specific cellular interactions triggers defense responses (6). Conceptually, these models would predict that immune cells are able to detect sudden internal disturbances and subsequently deploy some of their defensive arsenal. The extent of cellular perturbations that cells can recognize remains to be fully defined.

A hallmark of these models is the existence of sensors that indirectly recognize the presence of pathogens by monitoring the integrity of key host cell processes (6). In plants, one way that homeostasis sensors can trigger downstream signaling during microbial infection is through alteration of post-translational modification (PTM), including changes in protein phosphorylation, ubiquitination and glycosylation (7). PTMs facilitate rapid and dynamic control of protein function, and thus affect many cellular processes. Protein ubiquitination is a widespread and reversible PTM that occurs by covalent attachment of an ∼8. 5 kDa ubiquitin molecule to a target protein (8). Ubiquitin comprises up to 5% of total cellular protein content and is critical for regulation of many cellular processes including protein degradation, protein trafficking, signal transduction, apoptosis and innate immune signaling (9, 10). Therefore, tight control of cellular ubiquitin regulation is crucial for homeostasis, which is partially maintained through the ubiquitin-proteasome system (UPS) (11). However, in addition to its crucial role in protein degradation, ubiquitin is also a regulatory molecule, and most ubiquitination events in cells are associated with non-degradative regulatory functions (12). Control of protein ubiquitination is performed by the proteolytic action of deubiquitinase enzymes (DUBs), which remove or trim ubiquitin chains from modified proteins (13, 14). Another notable player that impacts ubiquitinated protein dynamics is the hexameric AAA ATPAse p97 (also called VCP/CDC48) (15, 16). Among other functions, p97 is involved in cellular quality control processes that result in the targeting and translocation of ubiquitinated proteins for remodeling, recycling or degradation (17–19). In immune cells, multiple components involved in the maintenance of the ubiquitin proteome have been associated with regulation of inflammatory responses, including p97 and many DUBs as exemplified by the activity of A20, DUBA, MYSM1 and OTULIN on immune signaling (20–23).

Regarding the importance of ubiquitination for many cellular processes, it is not surprising that pathogens have evolved to exploit or alter the ubiquitination system to promote infection (24). Recent proteomic studies have shown that infection can cause major alterations in the host ubiquitination/sumoylation (small ubiquitin-related modifier) profile. *Salmonella* Typhimurium infection of human colon cells causes substantial changes in the ubiquitination landscape, where 5 to 10% of all ubiquitination sites are differentially modified (25). Intoxication of cells with the LLO pore-forming toxin from *Listeria monocytogenes* or infection with live bacteria causes global alteration of the sumoylated proteome (26, 27). Similarly, viruses such as human immunodeficiency viruses (HIV-1 and −2) can hijack the host ubiquitin-proteasome system for their own benefit (28). These studies reinforce the idea that changes in protein PTMs, like ubiquitination, are a major target of manipulation by pathogens. It would therefore be advantageous for host cells to be able to detect perturbations in ubiquitin homeostasis in order to activate defense mechanisms.

Macrophages represent the first line of defense against invading pathogens and shape innate and adaptive immune responses (29). Here we investigate the effect of perturbing ubiquitin homeostasis, a common consequence of infection by pathogenic microbes, on the macrophage response to bacterial infection. Instead of starting with a complex infection model where multiple host signaling pathways are altered at once, we used a targeted chemical biology approach by using small-molecule compounds to perturb the cellular ubiquitin proteome *in vitro* (30–32). Using DUB inhibitors and a modulator of the p97 ATPase function, we show that macrophages respond to acute perturbation of cellular ubiquitin homeostasis by generating a robust and transient NOX2-dependent burst of reactive oxygen species (ROS) that promotes macrophage anti-infective capacity. Our results are consistent with a model whereby perturbation of the host ubiquitin landscape can be sensed by immune cells as a signal for activation of the macrophage anti-microbial arsenal.

## Results

### Perturbation of the ubiquitin proteome in macrophages results in accumulation of high-molecular weight ubiquitinated proteins

In order to globally perturb ubiquitin homeostasis in macrophages, we used a chemical biology approach. We previously characterized two structurally related DUB inhibitors, G9 (DUB^inh^) and C6 (DUB^inh^ C6), that cause global alteration of protein ubiquitination in macrophages at concentrations that preserve cell viability (Fig 1A) (30, 31). To better understand the impact of DUB^inh^ treatment on macrophage ubiquitination status, we compared the effect of DUB^inh^ with two other small molecules known to disrupt cellular ubiquitin homeostasis. RAW264. 7 cells, a murine macrophage-like cell line, were treated for 1h with DUB^inh^ or the p97/VCP modulator Eeyarestatin I (EerI) (32) or for 2h with the proteasome inhibitor MG132 (33). We observed that treatment of macrophages with DUB^inh^, EerI or MG132 resulted in notable increases in poly-ubiquitinated proteins, as expected (Fig 1B). Ubiquitin has seven lysine residues and an amino-terminal methionine that can be linked to other ubiquitin molecules. K48-ubiquitin chains target proteins for proteasomal degradation whereas K63-and M1-linkages have been associated with nonproteolytic signaling processes, including immune signaling (8, 34). All treatments described above induced an increase of K48-, K63-, and M1-specific ubiquitin linkages (Fig 1C). These results suggest that perturbation of the cellular ubiquitin proteome with small molecules results in accumulation of K48-ubiquitinated proteins, likely intended targets of the proteasome, as well as K63-and M1-modified proteins that could impact innate immune function. These data demonstrate that we can efficiently perturb cellular ubiquitin homeostasis in macrophages using a chemical biology approach.

**Figure 1:**
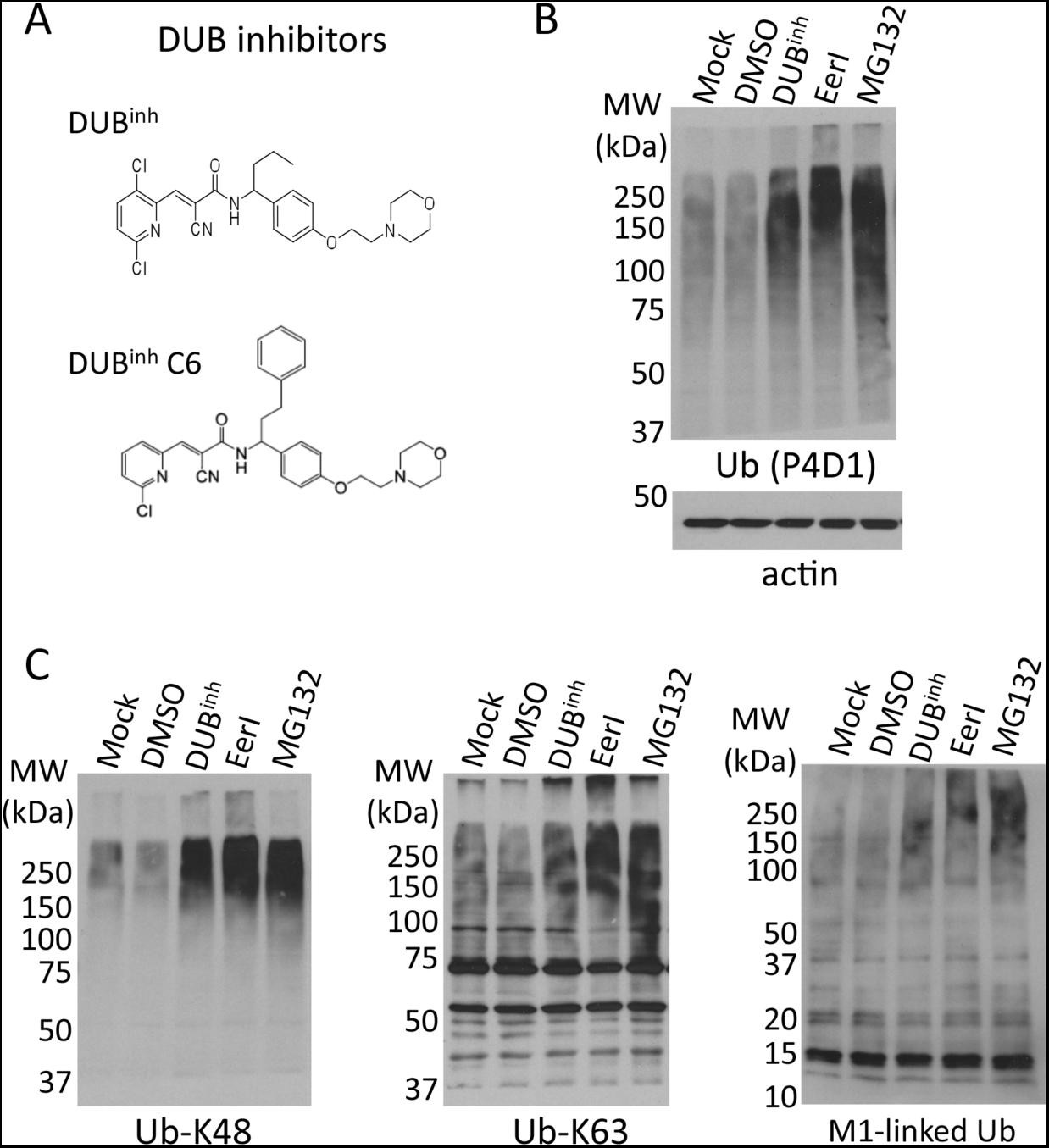
Molecules altering the ubiquitin proteome induce accumulation of ubiquitinated proteins. A). Schematic depicting the molecular structure of the DUB inhibitors G9 (DUB^inh^) and C6 (DUB^inh^ C6). B). Immunoblot of RAW264. 7 whole cell lysates showing accumulation of high-molecular weight ubiquitinated proteins following treatment with DUB^in^ (3. 5 µM, 1h), MG132 (5 µM, 2h) or EerI (10 µM, 1h). Immunoblot against **β-**actin was used as a loading control (bottom panel). C). Immunoblot of the same RAW264. 7 whole cell lysates probed with specific antibodies for K63-, K48-or M1-linked specific ubiquitin chains. Data are representative of at least three independent experiments.

### DUB^inh^ targets multiple DUBs in macrophages

The results shown above suggest that the small-molecule DUB^inh^ might target many cellular DUBs, causing a global alteration of the cellular ubiquitination profile. To further establish that multiple DUBs interact with DUB^inh^, we used a biotinylated version of this compound (DUB^inh^-biotin, Fig 2A) as an affinity reagent. As a control, we used a molecule with significantly reduced activity, ΔCN-biotin, lacking a key cyano group (30). RAW264. 7 cell lysates were incubated with DUB^inh^-biotin or ΔCN-biotin and protein complexes were isolated with streptavidin agarose beads. Silver staining of the precipitated proteins revealed that multiple proteins of various sizes specifically bind to DUB^inh^-biotin compared to ΔCN-biotin (Fig 2B). To further identify binding partners, we directly analyzed by immunoblot select DUBs involved in various cellular functions, including protein degradation (USP14) (35), endoplasmic reticulum-associated degradation (ERAD) (YOD1, Ataxin-3, USP25) (36–38) and innate immune signaling (MYSM1, DUBA, USP14) (21, 22, 39). We found that four out of the six interrogated DUBs, MYMS1, DUBA, Ataxin 3 and USP14, were associated with DUB^inh^-biotin, but to a lesser extent or not at all with ΔCN-biotin (Fig 2C). Our results are in agreement with a previous report suggesting that a variety of previously described DUB inhibitors have low selectivity and display activity against a wide range of DUBs (40). The ability of DUB^inh^ to bind multiple DUBs is consistent with the global increase in multiple ubiquitin linkage types (Fig 1C). Our results therefore supported the use of DUB^inh^ is a good chemical tool to assess the effects of general perturbation of ubiquitin homeostasis on the macrophage innate immune response.

### DUB inhibition induces ROS production in macrophages

Macrophages are important players of the innate immune system. We speculated that perturbation of macrophage ubiquitin homeostasis might trigger innate immune effector mechanisms. One main effector mechanism macrophages employ to kill invading pathogens and to promote immune signaling is the generation of reactive oxygen species (ROS). To determine whether DUB^inh^ treatment affects ROS production in macrophages, we loaded cells with the redox-sensitive fluorescent indicator chloromethyl-2’, 7’-dichlorodihydrofluorescein diacetate (CM-H_2_DCFDA) and treated them with vehicle control or DUB^inh^. By live cell flow cytometry analysis, ROS production was significantly increased in RAW264. 7 cells treated with DUB^inh^ compared to DMSO-treated cells (Fig 3A). Similar results were obtained using the DUB^inh^ C6 (Fig EV1A). Co-treatment of RAW264. 7 cells with DUB^inh^ and the ROS scavenger *N*-acetylcysteine (NAC) abolished ROS production (Fig. EV1B). Likewise, loading cells with reduced L-glutathione (GSH), a potent anti-oxidant, also significantly decreased ROS production in DUB^inh^-treated cells (Fig EV1B). We next investigated whether the effect of DUB inhibition on ROS generation also extended to the human monocyte cell lines THP-1 and U937. DUB inhibition using DUB^inh^ (Fig 3C–D) or DUB^inh^ C6 (Fig EV1C-D) also induced a burst of ROS early after treatment in these two human cell lines, suggesting that ROS generation may be a generalized response to ubiquitin homeostasis perturbation in monocytes. ROS are powerful effectors and signaling molecules; however, persistent ROS can be detrimental to cells (41). In order to understand the kinetics of ROS generation in response to DUB inhibition, we measured the ROS response at different times between 0. 5h and 5. 5h after treatment (Fig 3B). The increase in ROS production was observed as early as 0. 5h post-treatment with a peak response at 1. 5h after DUB^inh^. Strikingly, ROS levels decreased as early as 2. 5h post-treatment and returned to a level comparable to DMSO-treated macrophages by 4. 5h after treatment. Taken together, our results show that perturbation of ubiquitin homeostasis using DUB^inh^ induces a transient and robust ROS burst in mouse and human macrophage or monocyte cell lines.

**Figure 2:**
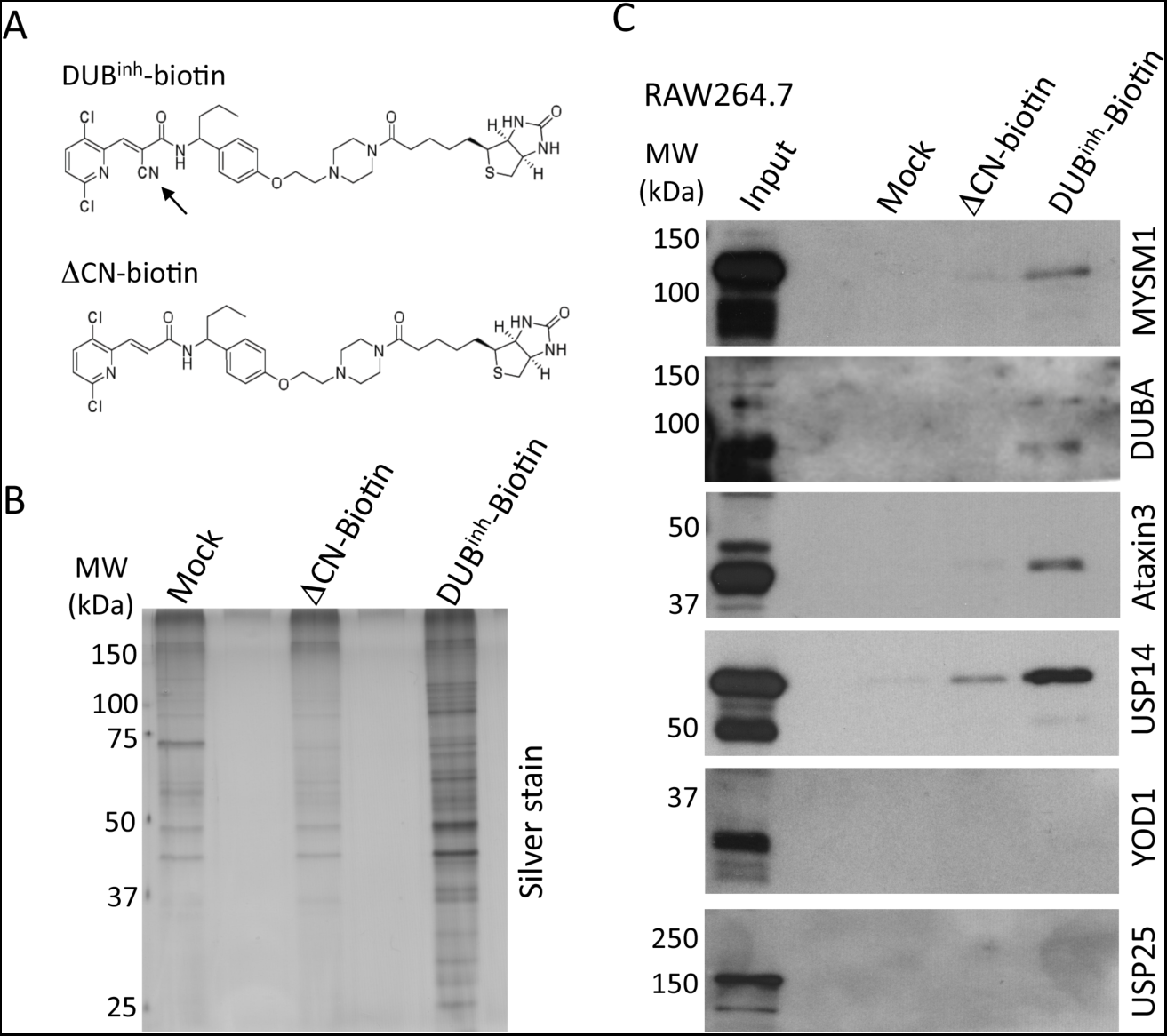
DUB inhibitor targets DUBs involved in multiple cellular functions. A). Structures of compounds DUB^inh^-Biotin and ΔCN-biotin. The arrow highlights the key cyano group. B). RAW264. 7 whole cell lysates were incubated with DUB^inh^-Biotin or ΔCN-biotin before immunoprecipitation using streptavidin-coated beads. The retained proteins were analyzed by SDS-PAGE and revealed by silver stain. C). Pull downs and corresponding inputs were immunoblotted for MYSM1, DUBA, Ataxin3, USP14, YOD1 and USP25. Data are representative of three independent experiments.

**Figure 3:**
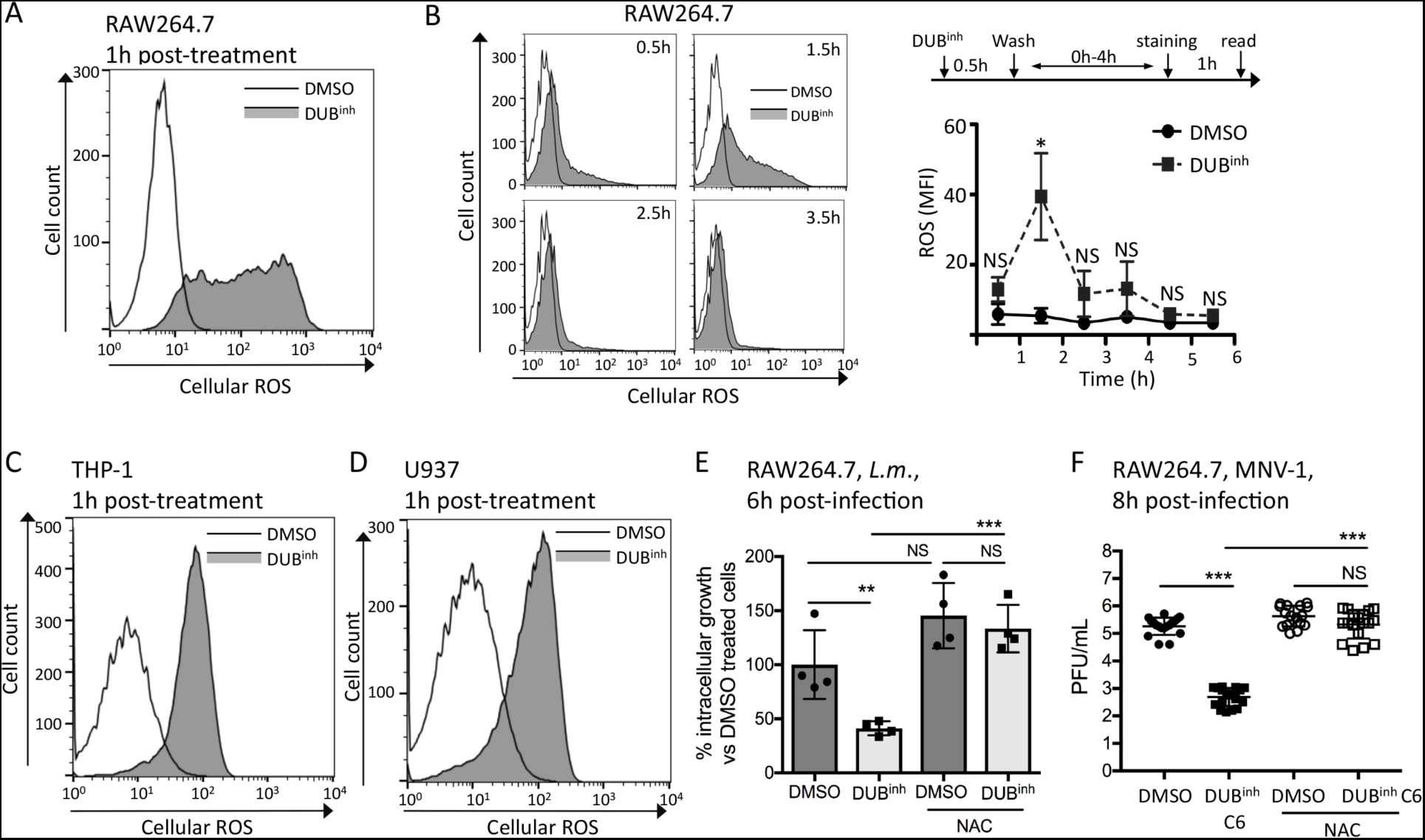
Inhibition of cellular DUBs in macrophages induces ROS generation. A). FACS analysis of RAW264. 7 cells treated with DUB^inh^ for 0. 5h before staining with 5 µM of the general ROS dye CM-H_2_DCFDA. FACS plots are representative of at least three independent experiments. B)Time course of ROS production in RAW264. 7 following treatment with 3. 5 µM DUB^inh^. The strategy used for the time course is depicted. Representative plots are shown for each time point and the indicated time represents the elapsed time after DUB^inh^ addition. Quantification of mean fluorescence intensity (MFI) was calculated from 3 independent experiments using FlowJo software. and D) FACS analysis of THP1 and U937 human monocytes treated with DUB^inh^ for 0. 5h before staining with 5 µM of CM-H_2_DCFDA. FACS plots are representative of three independent experiments. E) RAW264. 7 cells were incubated with 10 mM NAC or medium only for 0. 5h. DUB^inh^ (or equivalent volume of DMSO) was added on top at a final concentration of 3. 5 µM for 0. 5h. After incubation, the medium was removed, and cells were infected with *L. monocytogenes* at an MOI of 1 for 0. 5h. Following infection, cells were washed and new medium containing 10 µg/ml of gentamicin was added. Intracellular bacteria were enumerated at 6h post-infection. The data from four independent experiments performed in triplicate represent percent of intracellular *L. monocytogenes* growth compared to DMSO-only treated cells. F) RAW264. 7 cells were pre-treated with 10 mM NAC or medium in combination with 2. 5 µM DUB^inh^ C6 or equivalent volume of DMSO for 0. 5h before infection with MNV-1 at an MOI of 5 for 1h on ice. Cells were collected at 8h post-infection and viral titers were determined by plaque assay. Results are from three independent experiments. Data information: significant differences were calculated using one-way ANOVA and Dunnett’s post-test on the unmodified data (NS, not significant, * p < 0. 05, ** p < 0. 01, *** p < 0. 001).

ROS produced by phagocytes can have antimicrobial activity against a broad range of pathogens. We previously showed that DUB inhibition in RAW264. 7 cells resulted in lower pathogen burden using *Listeria monocytogenes* and murine norovirus (MNV-1), a surrogate for human norovirus, as model pathogens (30, 31). We chose these two distinct models of infection as both are intracellular pathogens that efficiently replicate within macrophages. To explore the relevance of ROS generation during DUB inhibition, we examined the effect of the ROS scavenger NAC and the anti-oxidant GSH on the antimicrobial activity of DUB^inh^ compounds. As shown before (30), treatment of RAW264. 7 cells with DUB^inh^ significantly reduced the number of *L. monocytogenes* intracellular colony forming units (CFU) at 6 hours post-infection (Fig 3E). However, addition of NAC or treatment with GSH during DUB inhibition significantly reduced the antimicrobial effect of DUB^inh^ against *L. monocytogenes*, consistent with a role for ROS generation as an antimicrobial effector (Fig 3E and EV1E). Likewise, addition of NAC or GSH abolished the effect of DUB^inh^ C6 on MNV-1 infection at 8 hours post-infection in RAW264. 7 cells (Fig 3F and EV1F). Collectively, our results suggest that perturbation of ubiquitin homeostasis using DUB^inh^ in macrophages promotes the generation of ROS that can effectively enhance antimicrobial effector function.

### Targeting function of the ubiquitin-dependent segregase p97 in macrophages induces ROS production

If perturbation of ubiquitin homeostasis is a global signal that promotes ROS production in macrophages, this response should also be induced by alteration of other regulators of the ubiquitin proteome. P97 is a gatekeeper for quality control of many different subcellular compartments, and its function relies on multiple ubiquitin-dependent cellular processes (16). As shown above (Fig 1B), altering p97 function using the EerI modulator caused a global shift in the ubiquitination profile of macrophages at a concentration that preserves cell viability. This observation correlates well with a previous study showing that p97 depletion in various cell lines also results in accumulation of poly-ubiquitinated proteins (42). Therefore, targeting p97 with EerI represents an alternative method for altering the ubiquitin proteome and perturbing ubiquitin homeostasis. RAW264. 7 cells were treated for 1h with EerI before staining with CM-H_2_DCFDA, followed by flow cytometry analysis. As with DUB^inh^ treatment, EerI caused a robust increase in ROS production (Fig 4A). Similar to the results shown above with DUB^inh^, the effect of EerI on macrophages was transient, as the ROS level decreased after 3h of treatment (Fig 4B). EerI treatment also induced ROS production in the human monocyte cells lines THP-1 and U937 (Fig 4C and D). To explore the potential effector function of ROS generated through EerI treatment, we used the MNV-1 infection model. As EerI directly prevents bacteria growth in broth culture, we could not use the *L. monocytogenes* infection model. Treatment of RAW264. 7 cells with 5 µM EerI resulted in a ∼2 log decrease in viral titer at 8h post-infection (Fig 4E). Therefore, as with DUB^inh^ treatment, disrupting p97 function also promoted ROS generation, reinforcing the idea that ubiquitin homeostasis perturbation is sensed by macrophages and results in activation of defense mechanisms.

**Figure 4:**
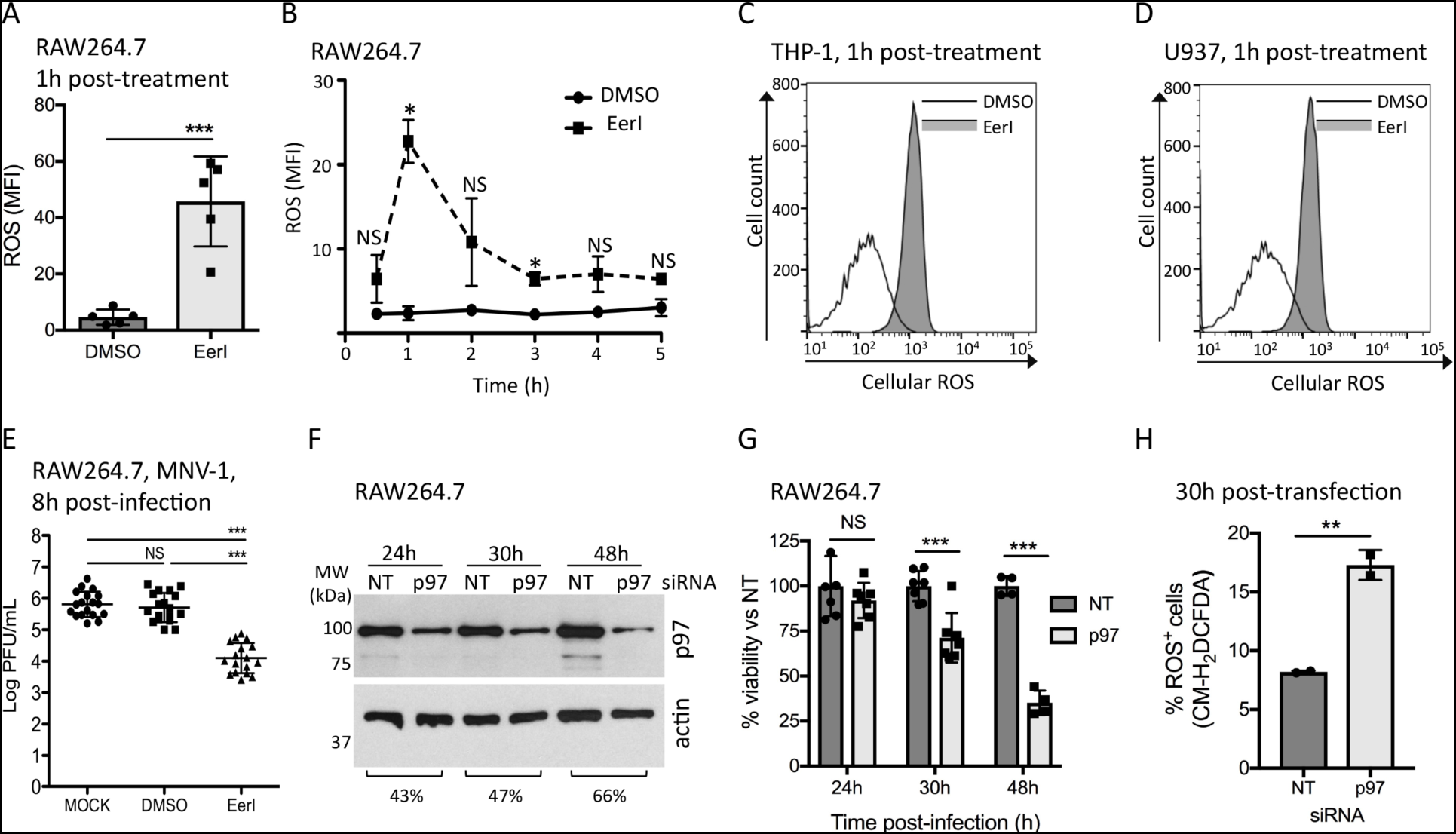
Perturbation of p97 activity in macrophages induces ROS generation. A). RAW264. 7 cells were treated with 10 µM EerI for 1h before staining with 5 µM of the general ROS dye CM-H_2_DCFDA (n = 5). B) Time course of ROS production in RAW264. 7 following treatment with 10 µM EerI for 1h. Quantification of mean fluorescence intensity (MFI) was calculated from 3 independent experiments using FlowJo software. and D) FACS analysis of THP1 and U937 human monocytes treated with 10 µM EerI for 1h before staining with 5 µM of CM-H_2_DCFDA. FACS plots are representative of three independent experiments. E). RAW264. 7 cells were treated with 10 µM EerI or equivalent volume of DMSO for 1h before infection with MNV-1 at an MOI of 5 for 1h on ice. Cells were collected at 8h post-infection and viral titers were determined by plaque assay. Results are from three independent experiments. F). RAW264. 7 were transfected with siRNA against p97 or a non-targeted control siRNA for the indicated period of time. Whole cells lysates were analyzed by immunoblotting against p97. The percentage of knock-down was calculated by band densitometry using actin to normalize the signal. G). At the indicated time point post-transfection, cells were washed and incubated for 90 minutes with the WST-1 cell viability reagent. Results represent the percent viability compared to the NT control transfected cells. Results are representative of two independent experiments performed in triplicate. H). At 30h post-transfection, cells were stained with 5 µM CM-H_2_DCFDA for 0. 5h. The percentage of cells stained for ROS (% ROS^+^ cells) were obtained from two independent experiments. Data information: significant differences were calculated using one-way ANOVA and Tukey’s post-test on the unmodified data (NS, not significant, * p < 0. 05, ** p < 0. 01, *** p < 0. 001).

As for all pharmacological approaches, there is a risk that compounds may induce off-target effects. To confirm the results obtained with EerI, we validated our hypothesis that perturbing p97 would trigger ROS production using a genetic approach. We used a pool of small interfering RNAs (siRNA) to knock-down p97 in murine macrophages. We first performed a time course after transfection to monitor knock-down efficiency and cell viability. As early as 24h post-transfection, we observed a ∼45% reduction in p97 protein levels compared to non-targeting (NT) control siRNA (Fig 4F). We reached ∼65% reduction of protein level by 48h post-transfection, however, notable loss of cell viability was also observed (Fig 4G), suggesting that p97 function is crucial for long-term macrophage survival. At 30h post-transfection, where we obtained suitable reduction in protein levels and limited loss of cell viability, we observed a 2-fold increase in the percentage of ROS-producing macrophages when transfected with p97 siRNA compared to NT control (Fig 4H). These results suggest that, as with DUB^inh^, disruption of p97 function might potentiate the antimicrobial capacity of macrophages through generation of ROS, consistent with our hypothesis that global ubiquitin perturbation is a key signal for activation of innate immune defense in macrophages.

### Perturbation of ubiquitin homeostasis causes increased ROS generation *ex vivo*

In order to assess the relevance of ubiquitin homeostasis perturbation for activation and signaling in primary immune cells, we tested the effect of DUB inhibition on ROS generation in primary murine macrophages. As observed using macrophage-like cell lines, DUB inhibition caused a burst of ROS in primary bone-marrow derived macrophages (pBMDM) (Fig 5A) as well as in immortalized bone marrow-derived macrophages (iBMDM) (Fig 5B). Prior activation with LPS and INF-γ was required for BMDM to elicit optimal ROS production. Similarly, treatment of peritoneal-resident macrophages with DUB^inh^ or p97 modulator EerI resulted in a significant increase in ROS generation (Fig 5C). We next used murine splenocytes, an *ex vivo* cell population that represents a more complex host immune environment. Splenocytes were treated *ex vivo* with either DUB^inh^ or EerI before staining with the ROS dye CM-H_2_DCDFA. Splenic macrophages were labeled using the F4/80 marker. We showed by flow cytometry that DUB^inh^ and EerI treatment also resulted in a significant increase in ROS production in this third physiologically relevant macrophage population (Fig 5D). These results suggest that ROS generation following perturbation of ubiquitin homeostasis is a broad and physiologically relevant response in both cultured and primary macrophages.

**Figure 5:**
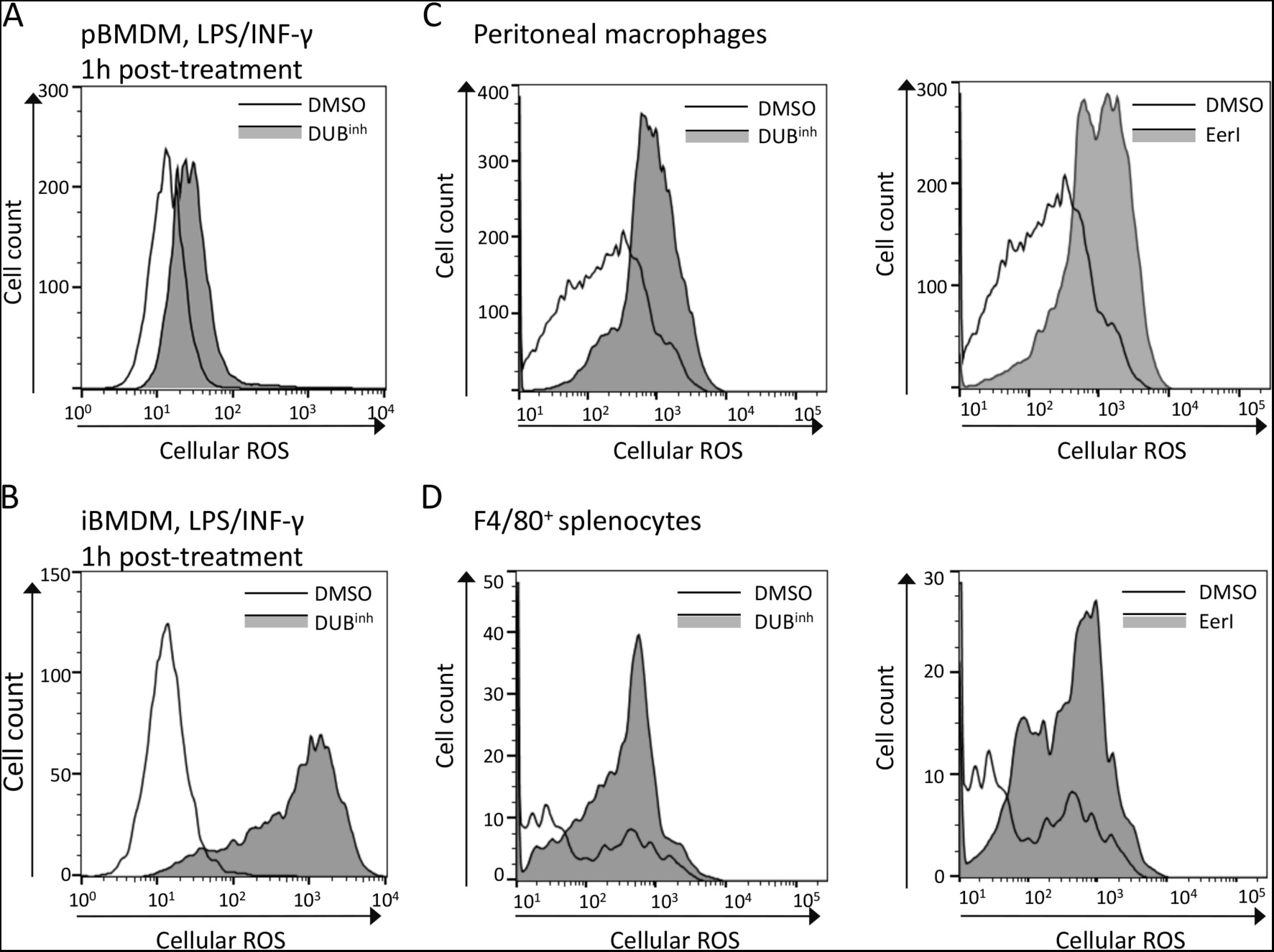
Perturbation of ubiquitin homeostasis induces ROS production in primary macrophages. A). and B) pBMDM (A) or iBMDM (B) incubated overnight with 100 ng/ml of LPS and INF-γ were treated for 0. 5h with 3. 5 µM DUB^inh^ before staining with 5 µM CM-H_2_DCFDA dye. C). Peritoneal macrophages were treated for 0. 5h with 1 µM DUB^inh^ or 1h with 10 µM EerI before staining with 1. 25 µM CM-H_2_DCFDA. D)Splenocytes were treated for 0. 5h with 3. 5 µM DUB^inh^ or 1h with 10 µM EerI before staining with 2 µM CM-H_2_DCFDA. Subsequent staining with F4/80^+^ antibody was performed to label the macrophage population. Data information: all FACS plots are representative of three independent experiments.

### ROS production upon perturbation of ubiquitin homeostasis in macrophages is independent of UPR activation

Perturbation of ubiquitin homeostasis can affect major cellular processes, including functions associated with the endoplasmic reticulum (ER), as protein ubiquitination is required for efficient removal and degradation of misfolded ER proteins (43). Perturbation of the balance between the folding capacity and the accumulation of unfolded proteins in the ER activates a signaling pathway termed the unfolded protein response (UPR) (44, 45). The UPR, mediated by three resident ER sensors, IRE1, PERK and ATF6, allows cells to adapt to ER stress and restore homeostasis. As the ER is essential for many cellular processes required for pathogen replication, it was proposed that this organelle can act as a surveillance platform to detect pathogen invasion and subsequently activate immune responses (46, 47). For example, activation of the UPR sensor IRE1 was recently associated with increased ROS production and bacterial killing in macrophages (48). We showed above that a consequence of DUB inhibition was the build-up of ubiquitinated proteins, a condition that can activate UPR signaling. Therefore, we probed the possible role of UPR signaling in ROS generation upon perturbation of ubiquitin homeostasis. RAW264. 7 macrophages were treated for 30 minutes with DUB^inh^ and samples were taken at 0. 5h, 2h and 6h after treatment. IRE1 activation was assessed by measuring the extent of *Xbp*1 transcript splicing, a direct target of the endonuclease domain of IRE1 (49). PERK activation was assessed by looking at induction of CHOP, encoded by a gene upregulated in a PERK-dependent manner (50). As a positive control for UPR activation, we used thapsigargin, a potent inhibitor of the sarco-endoplasmic reticulum Ca^2+^-ATPase (SERCA pump), and tunicamycin, a protein glycosylation inhibitor (45). Thapsigargin and tunicamycin treatment induced CHOP expression and *Xbp1* splicing, as expected (Fig 6A–B). However, no activation of either IRE1 or PERK pathways was detected in cells treated with DUB^inh^ (Fig 6A-B). To confirm that IRE1 activation was not involved in generation of antimicrobial ROS upon DUB inhibition, we used the IRE1 inhibitor 4µ8c (51). Pre-treatment of cells with 4µ8c did not rescue *L. monocytogenes* growth inhibition in RAW264. 7 cells upon DUB inhibition (Fig 6C). Similarly, inhibition of PERK kinase activity using GSK 2606414 (GSK-PERK) (52) had no effect on DUB^inh^-induced antimicrobial activity against *L. monocytogenes* (Fig 6D). To more broadly test the possible role of UPR signaling in DUB^inh^-mediated ROS production, we treated cells with tauroursodeoxycholic acid (TUDCA), a chemical chaperone that ameliorates ER stress (53). RAW264. 7 cells produced similar levels of ROS after DUB^inh^ treatment in the presence or absence of TUDCA (Fig 6E). These results suggest that ROS generation following ubiquitin homeostasis perturbation by DUB^inh^ does not rely on the unfolded protein response.

**Figure 6:**
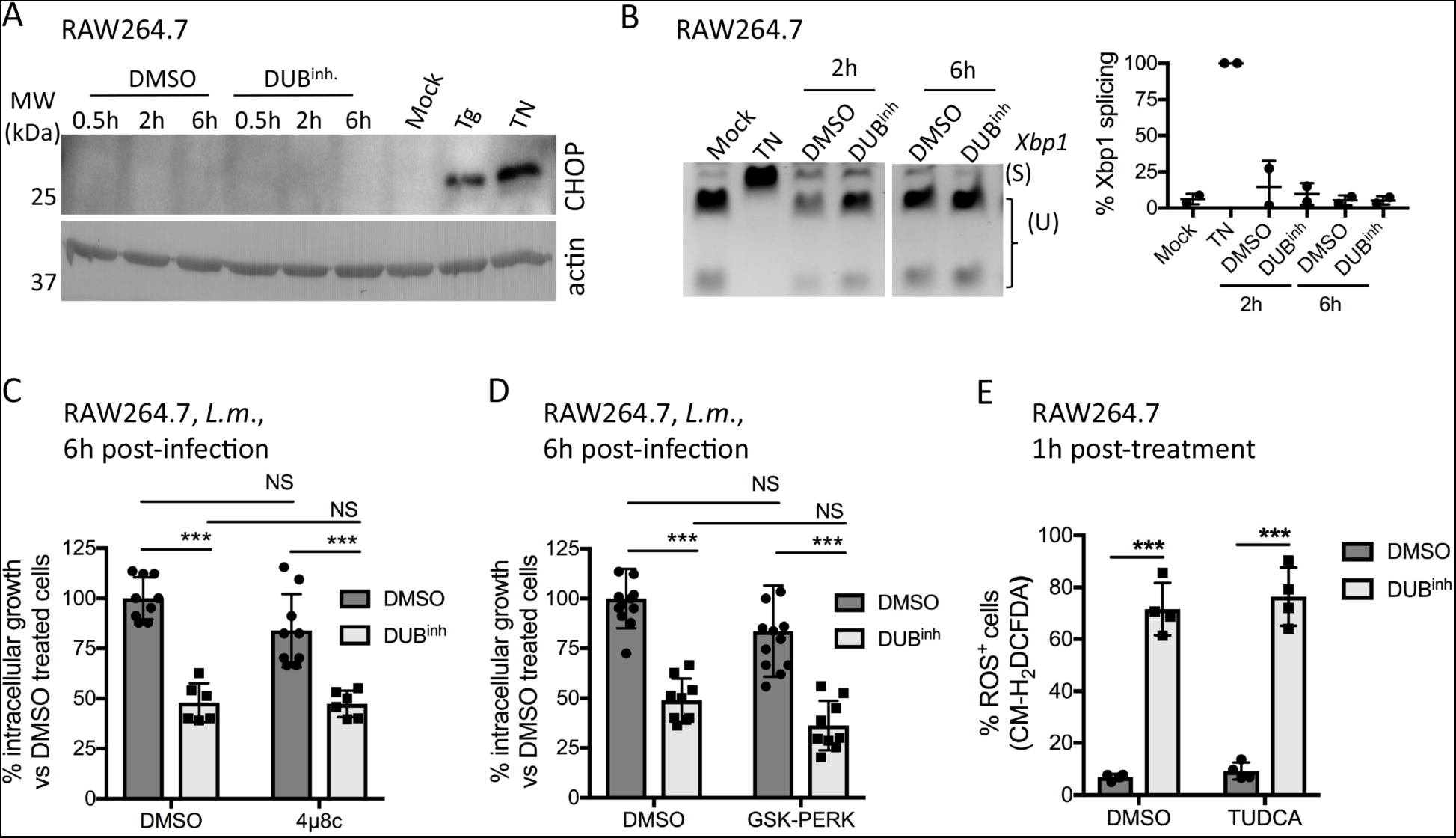
DUB^inh^ induces ROS generation independently of UPR signaling. A). RAW264. 7 cells were treated for 0. 5h with 3. 5 µM DUB^inh^. Following treatment, cells were washed and subsequently incubated for the indicated period of time in fresh medium. As control, cells were treated with 10 µM thapsigargin or 10 µg/ml of tunicamycin for 4h. Whole cells lysates were used for immunoblotting against CHOP or actin as a loading control. B) RAW264. 7 were treated as above. PCR was performed to amplify the *Xbp1* mRNA and products were digested with the Pst1 endonuclease. The unspliced (U) form of *Xbp1* contains a Pst1 cleavage site, which will generate two smaller fragments compared to the *Xbp1* spliced (S) form. The percentage of *Xbp1* splicing was calculated by band densitometry as follow: *Xbp1*(S) / [*Xbp1*(U)+*Xbp1*(S)]. Results for A) and B) are from two independent experiments. C) and D) RAW264. 7 cells were incubated for 0. 5h with 50 µM 4µ8C (C) or 50 µM GSK-PERK D) (D). DUB^inh^ (or equivalent volume of DMSO) was added on top at a final concentration of 3. 5 µM for 0. 5h. After incubation, the medium was removed, and cells were infected with *L. monocytogenes* at an MOI of 1 for 0. 5h. Following infection, cells were washed and new medium containing 10 µg/ml of gentamicin was added. Intracellular bacteria were enumerated at 6h post-infection. The data represent percent of intracellular *L. monocytogenes* growth compared to DMSO-only treated cells. Results were obtained from three independent experiments performed in triplicate. E) RAW264. 7 cells were incubated with 300 µM TUDCA or medium only for 1h. DUB^inh^ (or equivalent volume of DMSO) was added on top at a final concentration of 3. 5 µM for 0. 5h before staining for ROS detection. The results from four experiments represent the percentage of cells stained for ROS (% ROS^+^ cells). Data information: significant differences were calculated using two-tailed student’s t-test or one-way ANOVA and Tukey’s post-test on the unmodified data (NS, not significant, *** p < 0. 001).

### Inhibition of proteasome function is not sufficient to induce ROS generation in macrophages

One well-known function of p97 is to trigger protein extraction from membranes or complexes in order to facilitate degradation by the proteasome (16). Moreover, DUB function is integral to the ubiquitin-proteasome system, and we showed above that inhibition of proteasome function also resulted in global perturbation of cellular ubiquitination (Fig 1B). Consequently, we next investigated the effect of proteasome inhibition on ROS production by macrophages. RAW264. 7 cells were treated for 1h with MG132 or epoxomicin, two commonly used proteasome inhibitors, and ROS production was measured as before. In contrast to the results obtained with the DUB^inh^ and EerI, no increase in ROS generation was observed (Fig 7A). It is important to note that EerI and other well-studied DUB inhibitors have been shown to have no direct effect on proteasome proteolytic activity (54). Also, p97 inhibition resulted in distinct alterations of the ubiquitin-modified proteome compared to proteasome inhibition, despite their similar effect on global ubiquitin homeostasis (12). These observations reveal that different insults to the cellular ubiquitin proteome may stimulate distinct cellular responses. Accordingly, our findings raise the possibility that perturbation of ubiquitin homeostasis in macrophages by altering DUBs or p97 activity results in a different signal compared to direct inhibition of proteasome function.

**Figure 7:**
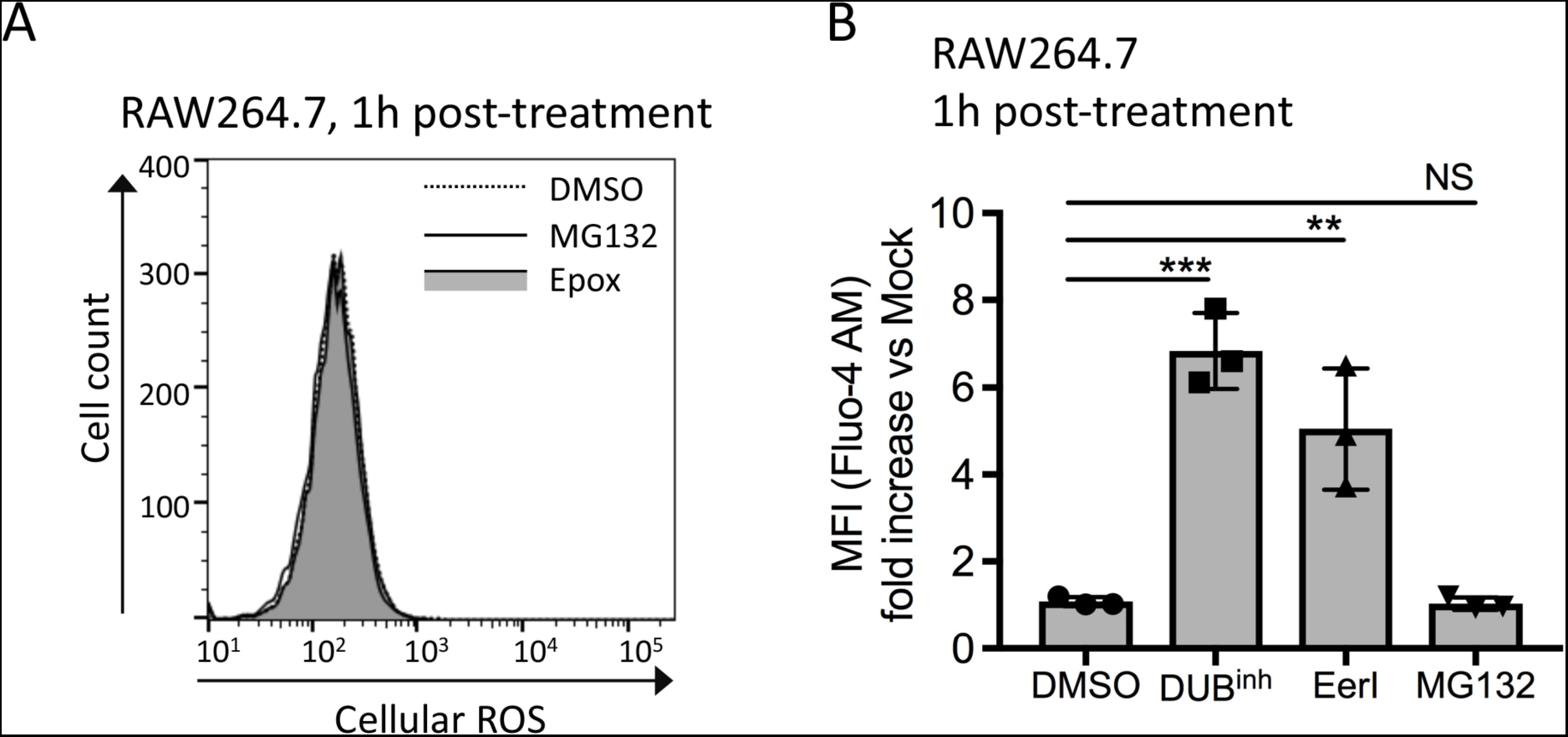
Specific perturbation of ubiquitin homeostasis alters intracellular calciumlevels. A). RAW264. 7 cells were treated with 5 µM MG132 or 0. 5 µM epoxomicin for 2h before staining for ROS. FACS plots are representative of three independent experiments. B). RAW264. 7 cells were treated for 2h with 5 µM MG132, 1h with 10 µM EerI or 0. 5h with 3. 5 µM DUB^inh^ or equivalent volume of DMSO before staining for 0. 5h with 1 µM Fluo-4 AM calcium indicator. Results represent fold increase in mean fluorescence intensity (MFI) compared to untreated cells. Results are from three independent experiments. Data information: significant differences were calculated using one-way ANOVA with Tukey’s multiple comparison test (NS, not significant, ** p < 0. 01, *** p < 0. 001).

### Alteration of ubiquitin homeostasis using DUB^inh^ or EerI increases intracellular calcium

Recent evidence links alteration of protein ubiquitination with calcium flux. For example, expression of a ubiquitin mutant containing a K6W substitution, which can be conjugated to protein but is proteolytically incompetent and causes accumulation of ubiquitin conjugates, resulted in a 4-fold increase in intracellular calcium level in the ocular lens (55). Calcium ions are important second messengers for multiple signaling pathways, including those regulating ROS production (56). To assess the effect of DUB inhibition and perturbation of p97 function on intracellular levels of calcium, we used the Fluo-4 AM calcium indicator, which exhibits increased fluorescence upon calcium binding and can be quantified by flow cytometry. Staining of RAW264. 7 treated with DUB^inh^ using Fluo-4 AM showed a ∼6-fold increase in mean fluorescence intensity (Fig 7B). Similar results were obtained when macrophages were treated with EerI (Fig 7B). As mentioned previously, divergent insults to the ubiquitin proteome might result in distinct cellular responses. Thus, we next examined the effect of proteasome inhibition on intracellular calcium. Notably, we observed no change in calcium levels upon MG132 treatment (Fig 7B). Our results demonstrate that perturbation of ubiquitin homeostasis by altering DUBs or p97 function, but not through proteasome inhibition, stimulates an increase in intracellular calcium levels.

### Perturbation of DUBs or p97 function results in NOX2-dependent ROS production

The NADPH phagocyte oxidase (NOX2) is a major source of ROS in macrophages. The NOX2 core is composed of membrane-bound p22^phox^ and gp91^phox^, the cytochrome b558 unit. NOX2 complex activation is regulated by multiple cytosolic subunits, including p40^phox^, p47^phox^, p67^phox^ and the Rac GTPase (57). The function and localization of some of these subunits are controlled by PTMs, emphasizing the complexity regulating NOX2 activation. We investigated the role of the NOX2 complex in DUB^inh^-mediated ROS production. iBMDM isolated from WT or *gp91*^*phox-/y*^ mice, which lack a functional NOX2 complex, were treated with DUB^inh^ for 0. 5h and then assayed for cellular ROS. A ∼6-fold decrease in fluorescence was observed between WT and *gp91*^*phox-/y*^ iBMDM (Fig 8A), suggesting an active role for NOX2 in generating ROS in response to DUB^inh^. We also looked at whole cell extracts from RAW264. 7 cells or iBMDM after treatment with DUB^inh^ and observed no consistent change in gp91^phox^ expression levels (Fig 8B). *Gp91*^*phox-/y*^ iBMDM still produced low levels of ROS in response to DUB^inh^, suggesting a minor secondary source of ROS production. As macrophages possess other NADPH oxidase enzymes, we tested the effect of NOX1 or NOX4 loss of function on ROS generation after DUB^inh^ treatment. Using primary BMDM isolated from *Nox1*^-/-^ or *Nox4*^-/-^ mice, we observed that these two NADPH oxidase enzymes are dispensable for ROS generation following DUB^inh^ treatment (Fig 8C). EerI-induced ROS in macrophages was also dependent on NOX2 activity as a ∼4 fold-decrease was observed in iBMDM lacking the gp91^phox^ subunit compared to WT cells (Fig 8D). If NOX2 is a major effector protein sensing perturbation of ubiquitin homeostasis, we would expect the anti-microbial activity of DUB^inh^ compound to be reduced or abolished in cells lacking this NADPH oxidase complex. Accordingly, we tested the effect of DUB inhibition on *L. monocytogenes* growth in macrophages lacking gp91^phox^. As shown in Figure 8E, DUB inhibition in WT iBMDM resulted in a significant decrease of *L. monocytogenes* growth at 6h post-infection. However, no significant difference was observed for *L. monocytogenes* grown in *gp91*^*phox-/y*^ iBMDM treated with DUB^inh^ compared to the DMSO-vehicle control, consistent with our hypothesis. Taken together, our data show that upon perturbation of ubiquitin homeostasis, macrophages generate a burst of ROS that is largely dependent on the NOX2 phagocyte oxidase.

**Figure 8:**
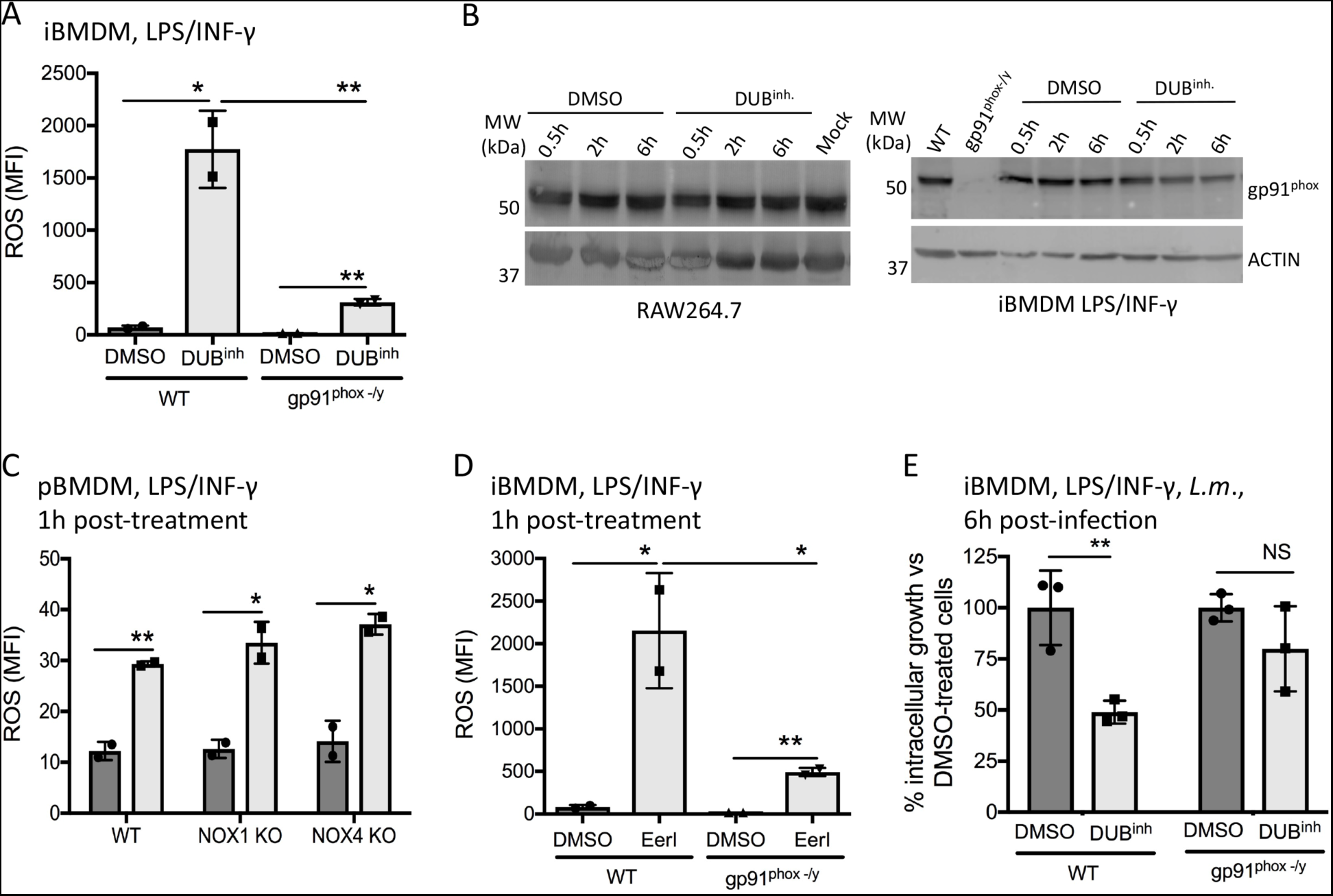
Perturbation of ubiquitin homeostasis induces NADPH phagocyte oxidase-dependent ROS generation in macrophages. A). WT and gp91^phox -/y^ iBMDM incubated overnight with 100 ng/ml of LPS and INF-γ were treated for 0. 5h with 3. 5 µM DUB^inh^ or equivalent volume of DMSO before staining for 0. 5h with 5 µM CM-H_2_DCFDA. Quantification of mean fluorescence intensity (MFI) from two independent experiments was calculated using FlowJo software. B). RAW264. 7 cells or WT and gp91^phox -/y^ iBMDM incubated overnight with 100 ng/ml of LPS and INF-γ were treated for 0. 5h with 3. 5 µM DUB^inh^. Following treatment, cells were washed and subsequently incubated for the indicated period of time in fresh medium. Whole cells lysates were used for immunoblotting against gp91^phox^ or β-actin as a loading control. Results are from two independent experiments. C). pBMDM isolated from WT, *Nox*^*-/-*^ (NOX1 KO) or *Nox4*^*-/-*^ (NOX4 KO) mice were incubated overnight with 100 ng/ml of LPS and INF-γ and treated for 0. 5h with 3. 5 µM DUB^inh^ or equivalent volume of DMSO before staining for 0. 5h with 5 µM CM-H_2_DCFDA. Results are from two independent experiments. D). WT and gp91^phox -/y^ iBMDM incubated overnight with 100 ng/ml of LPS and INF-γ were treated for 1h with 10 µM EerI or equivalent volume of DMSO before staining for 0. 5h with 5 µM CM-H_2_DCFDA. Results are from two independent experiments. E). WT and gp91^phox -/y^ iBMDM incubated overnight with 100 ng/ml of LPS and INF-γ were treated for 0. 5h with 3. 5 µM DUB^inh^. After incubation, the medium was removed, and cells were infected with *L. monocytogenes* at an MOI of 1 for 0. 5h. Following infection, cells were washed and new medium containing 10 µg/ml of gentamicin was added. Intracellular bacteria were enumerated at 6h post-infection. The data represent percent of intracellular *L. monocytogenes* growth compared to DMSO-only treated cells. Results are from three independent experiments. Data information: significant differences were calculated using two-tailed student’s t-test or one-way ANOVA and Tukey’s post-test (NS, not significant, * p < 0. 05, ** p < 0. 01).

## Discussion

Emerging evidence indicates that cells recognize sudden perturbations of normal cellular functions as a signal to activate innate immune defenses (2, 5). Here, we propose a new homeostasis-associated molecular pattern, perturbation of ubiquitin homeostasis, that can be sensed by macrophages as a trigger for antimicrobial activity. As infection itself can change the ubiquitin landscape of cells, and multiple bacterial and viral pathogen genomes encode proteins that hijack the host ubiquitination machinery, it is conceivable that cells possess mechanisms to detect these changes. Therefore, we used a chemical biology approach to cause global alterations in the ubiquitin proteome in order to study specific macrophage responses. Treatment of macrophages with either a DUB inhibitor or the p97 modulator EerI resulted in increased ubiquitinated proteins, including K48-, K63-and M1-specific linkages, validating our approach. Notably, we observed that perturbation of ubiquitin homeostasis in human and murine macrophage cell lines and in primary macrophages results in a robust but transient NOX2-dependent generation of reactive oxygen species.

In order to establish conditions favorable for survival and proliferation in a host, pathogens employ multiple strategies, including modification of host signaling pathways (58, 59). Many pathogens inject effector proteins that disrupt the cytoskeleton, whereas other pathogens use or block the function of the host translation machinery or hijack the protein synthesis and folding machineries of the ER in order to promote their own survival. It is therefore crucial for the cell to guard these fundamental processes as they are often essential for cell survival and function. This concept of a “guard model” has been well studied in plant immunity, where PTM of guard proteins is a common sensing mechanism, highlighting the importance of PTMs for this system. Recently, a molecular basis for this concept was established in mammalian cells using macrophages. The pyrin inflammasome is activated upon loss of pyrin phosphorylation resulting from RhoA small GTPase inactivation by pathogens, which allows the release of the inhibitory protein 14-3-3 and inflammasome activation (60). Therefore, similar to the guard proteins in plant immunity, the mammalian pyrin inflammasome acts as a global sensor that is modulated by PTMs. However, the possible guard sensors identified so far in mammalian cells are limited and often speculative, therefore more work is required to understand their function and the signals triggering their response.

Here we showed that perturbation of ubiquitin homeostasis in macrophages induced ROS generation via the NOX2 complex. In phagocytic cells, the best-known function of NOX2-dependent ROS is antimicrobial activity (61). Accordingly, we showed that ROS generated upon DUB inhibition results in an increased capacity of macrophages to control viral and bacterial intracellular infections. However, besides its antimicrobial function, NADPH-generated ROS can also act as second messenger important for signaling (62). NADPH oxidase activity is associated with the activation of other well-known macrophage effector functions, including autophagy and inflammasome activation (63). At an organismal level, NADPH-generated ROS are also key signaling molecules for immune cell recruitment at wound or infection sites and are also involved in modulation of adaptive immune responses (62, 64). Thus, generation of ROS by macrophages following homeostasis perturbation might be a strategic way to target multiple effector mechanisms that will potentially allow protection against a broad spectrum of pathogens.

Regulation of NOX2 NADPH oxidase activity is multi-layered and this complex requires at least five subunits to be functional; the heterodimer transmembrane proteins gp91^phox^ and p22^phox^, the three-cytosolic subunits p47^phox^, p40^phox^ and p67^phox^ and RAC, a small regulatory GTPase (57). Activation of the NOX2 complex is controlled through alterations of the cytosolic subunits, especially through phosphorylation events. Besides phosphorylation, other PTMs play active roles in the control of NOX enzymes, including ubiquitination and sumoylation, even though limited information is available to date. Nevertheless, the NADPH oxidases NOX2, NOX4 and NOX5 were shown to be ubiquitinated or sumoylated, modifications that directly impact ROS generation (65–67). RAC1, a critical component for NOX2 activation, is also modified by ubiquitination (68). Similarly, the p47^phox^ subunit of the NOX2 complex can be ubiquitinated in macrophages. However, the function of this PTM for p47^phox^ remains unknown (69). As post-translational modifications are a key control point for innate immune signaling and a way to sense pathogen-induced perturbations, it is tempting to speculate that alterations in ubiquitin homeostasis affect the function of at least one regulatory component of the NOX2 complex. However, further study will be required to specifically uncover the direct or indirect effect of ubiquitin perturbation on each subunit and to assess the possible role of the NOX2 complex as a homeostasis guard sensor.

Perturbation of ubiquitin homeostasis can be associated with an accumulation of unfolded or non-native proteins in the ER, a condition leading to ER stress and activation of the UPR (44, 45). Signaling downstream of UPR sensors has been shown to affect key innate immune responses, including NF-κB activation, cytokine production and ROS generation (59, 70-72). The ER itself is also an important source of cellular ROS as its lumen is the most oxidizing environment in the cell, an attribute required to support protein folding. However, despite an established link between perturbation of the ubiquitin proteome and induction of ER stress, we were able to show here that DUB inhibition was not associated with UPR activation in macrophages. Moreover, we showed that alleviation of ER stress using a chemical chaperone had no effect on DUB inhibition-induced ROS, suggesting that DUB inhibition-induced ROS generation does not rely on activation of the UPR.

Recent proteomic studies have highlighted major differences in the repertoire of changes to the cellular ubiquitin proteome that depend on the trigger used to perturb ubiquitin dynamics. It was shown that proteasome and p97 inhibition, both resulting in increased global poly-ubiquitinated protein accumulation, elicit unique and distinct alterations to the ubiquitin-modified proteome (12). Accordingly, proteasome inhibition was shown to mainly stabilize newly synthesized proteins, suggesting that a major proteasome function is targeting defective translation products for degradation (73). As ubiquitination events mainly control protein function in a non-degradative way, it is tempting to hypothesize that altering p97 or DUB activity would have a broader impact on cells by targeting multiple regulatory proteins. Indeed, we showed here that proteasome inhibitors failed to induce ROS generation in macrophages despite their ability to alter ubiquitin homeostasis in cells. Moreover, DUB inhibition and perturbation of p97 function, but not proteasome inhibition, resulted in increased intracellular calcium levels. Importantly, these observations suggest a clear distinction in macrophage responses to different signals that affect ubiquitin homeostasis. It is interesting to note that a similar intracellular calcium increase was observed in lens cells overexpressing a mutant form of ubiquitin that caused accumulation of ubiquitinated proteins without directly affecting proteasome function (55). Calcium is a powerful signaling molecule and is involved in activation of many signaling pathways, including those leading to NOX2 activation (56). It is thus possible that ubiquitin homeostasis perturbation indirectly triggers ROS generation by increasing calcium intracellular levels, but further study will be needed to specifically determine the interplay between these two signals.

By using compounds targeting two distinct cellular processes, we show that alteration of ubiquitin status is a trigger for macrophages to activate innate immune pathways. Uncovering cellular processes that are altered during infection is crucial for a full understanding of the activation mechanisms necessary for innate immune responses. Moreover, this knowledge may also be applied to better understand a subset of autoimmune diseases. Substantial alterations in cellular homeostasis have been linked with many conditions associated with increased inflammation, including obesity, hypertension, type 2 diabetes, and atherosclerosis (74). Therefore, our results lead us to propose global ubiquitin perturbation as a potent new signal that triggers an acute defense response in macrophages with the potential to control a broad spectrum of pathogenic infections.

## Materials and methods

### Antibodies and reagents

DUB^inh^-related compounds were dissolved in dimethyl sulfoxide (DMSO), aliquoted and stored at −80^°^C. The two DUB inhibitors, DUB^inh^ (previously called G9 or compound 9) and DUB^inh^ C6, used in this study have been described elsewhere (30, 31). The DUB inhibitor G9 is used extensively in this study and therefore is called DUB^inh^ throughout the manuscript. The DUB inhibitor C6 was used to confirm results and is labelled as DUB^inh^ C6 in the text. DUB^inh^-biotin and ΔCN-biotin were synthesized by the University of Michigan Vahlteich Medicinal Chemistry Core. Full experimental details around the synthesis of DUB^inh^, DUB^inh^-biotin, and ΔCN-biotin are given in the Appendix supplementary methods and figures as Figures S1 to S3, respectively. The library numbers for each compound from the University of Michigan (CCG) database are CCG-257343 for DUB^inh^, CCG-257202 for DUB^inh^-biotin and CCG-257203 for ΔCN-biotin. Eeyarestatin I, GSK2606414 (GSK-PERK), 4µ8C, Tauroursodeoxycholic acid (TUDCA) and thapsigargin were purchased from Millipore. MG132 was purchased from Selleckhem and epoxomicin from Cayman Chemical. The general ROS indicator CM-H_2_DCFDA and the calcium indicator Fluo-4 AM were from ThermoFisher, whereas reduced L-glutathione, WST-1 cell viability reagent and N-acetylcysteine (NAC) were purchased from Sigma. Bacterial lipopolysaccharide (LPS) was purchased from Abcam and recombinant murine interferon-γ from Peprotech. The primary antibodies used for immunoblotting were anti-ubiquitin, clone P4D1 (sc-8017), anti-USP14 (sc-100630), anti-USP25 (sc-398414) and anti-gp91^phox^ (sc-130543) from Santa Cruz, anti-YOD1 (ARP67915) from Aviva Systems Biology, anti-Otud5 (DUBA, 21002-1-AP) from ProteinTech, anti-MYSM1 (ab193081) and anti-Ataxin3 (ab175262) from Abcam, anti-ubiquitin lys48-specific (05-1307) and anti-linear ubiquitin clone LUB9 (MABS451) from EDM Millipore, anti-ubiquitin k63-specific (5621) and anti-CHOP (2895) from Cell Signaling, anti-actin clone ACTN05 (MS-1295) and anti-p97/VCP (MA3-004) from TermoFisher. All HRP-conjugated secondary antibodies were from Jackson ImmunoResearch. The silver stain was done using the Pierce silver stain kit (ThermoFisher).

### Cells, mouse strains, bacterial strains and viruses

Wild-type C57BL/6 Mice were housed in specific pathogen-free facilities, maintained by the Unit for Lab Animal Medicine of the University of Michigan. This study was carried out in accordance with the recommendations in the guide for the care and use of laboratory animals of the National Institutes of Health and the protocol was approved by the committee on the care and use of animals of the University of Michigan. NOX2-deficient (*cybb*^-/y^), *Nox1*-and *Nox4*-deficient mice and age-matched wild-type C57BL/6 mice were purchased from Jackson Laboratories. Male and female mice aged 8-12 weeks were used. Animal experiments were not randomized or blinded. Bone-marrow derived macrophages (BMDM) were prepared by flushing mouse femurs in Dulbecco’s modified Eagle’s medium (DMEM) medium. Differentiation was done by incubating cells in BMDM media containing 50% DMEM, 30% L929-conditioned medium, 20% heat-inactivated fetal bovine serum (FBS), 5% L-Glutamine, 0. 05% β mercaptoethanol and 100 units/ml of penicillin-streptomycin (Pen/strep) for six days. To generate immortalized BMDM (iBMDM), freshly harvested bone-marrow cells were transduced with the J2 retrovirus and differentiated in macrophages as above (75). iBMDM were maintained in DMEM medium containing 10% FBS, 5% L-Glutamine, 1% non-essential amino acids (NEAA) and 1% HEPES. Peritoneal macrophages were harvested after washing the peritoneal space twice with 3-ml of cold PBS. Splenocytes were obtained by passing the spleen through a 70 µM cell strainer. Red blood cells were removed by incubation of splenocytes at room temperature for 2 minutes in lysis buffer (155 mM ammonium chloride, 12 mM sodium chloride and 0. 1 mM EDTA pH8). Splenocytes were stained with 1 µg of phycoerythrin (PE) rat anti-mouse F4/80 antibody (BD biosciences #565410) in order to label the macrophage population. RAW264. 7 cells, THP-1 cells and U937 cells were originally obtained from American Type Culture Collection (ATCC). All cell lines were originally authenticated by ATCC and were not further tested in our laboratory. RAW264. 7 cells were grown in DMEM supplemented with 10% FBS, 5% L-Glutamine, 1% NEAA, 1% HEPES and Pen/Strep, whereas THP-1 and U937 cells were grown in RPMI 1640 medium supplemented with 10% FBS, 5% L-Glutamine, 1% NEAA, 1% HEPES, 0. 05% β-mercaptoethanol and Pen/Strep. All cells were incubated at 37^°^C with 5% CO_2_. Cells were tested negative for mycoplasma contamination. *L. monocytogenes* 10403S was grown in brain hearth infusion (BHI) broth statically at 30^°^C overnight. The plaque-purified MNV-1 strain (GV/MNV1/2002/USA) MNV1. CW3 (referred as MNV-1 in the text) was used at passage 6 for all experiments (76).

### Protein extraction, pull down and immunoblotting

For pull-down experiments using DUB^inh^-biotin probe, RAW264. 7 cells were seeded at a density of 7 × 10^6^ cells per 100 mm dish and allowed to adhere overnight. Cells were washed twice with Dulbecco’s phosphate-buffered saline (PBS) and lysed in ice-cold cell lysis buffer (1% NP-40, 150 mM NaCl, 50 mM Tris-Hcl pH8, 1mM EDTA pH8, 0. 1% SDS, 0. 5% sodium deoxycholate, 1X Roche protease inhibitor). After 15 minutes incubation on ice, cells were briefly sonicated and split into 4 tubes. The input tube was diluted in 4X concentrated sample buffer (BioRad) containing β-mercaptoethanol and denatured for 10 minutes at 95^°^C. The remaining tubes were either left untreated (mock sample) or incubated at 37^°^C for 30 minutes with 20 µM DUB^inh^-biotin or ΔCN-biotin. Pre-washed streptavidin agarose resin (Pierce) was added directly to the lysate and tubes were incubated with agitation for 2h at 4^°^C. Agarose beads were washed 6 times with lysis buffer and resuspended in 75 µl of 1 × sample buffer. To obtain whole-cell lysates, RAW264. 7 cells or iBMDM cells were seeded in 6-well plates at an initial density of 6 × 10^6^ cells per plate. iBMDM were activated overnight with 100 ng/ml of LPS and interferon-γ. After treatment described in the specific figure legends, cells were lysed in a cell lysis buffer (10 mM Tris-HCl pH8, 150 mM NaCl, 1 % NP-40, 10 mM EDTA pH8, 1 mM DTT and 1X Roche protease inhibitors), incubated on ice for 15 minutes, quickly sonicated and diluted in 4X sample buffer. Samples were separated by SDS-PAGE and transferred to polyvinyldene fluoride membrane (PVDF, Millipore). Immunoblotting was performed according to the antibody manufacturers’ instructions.

### *L. monocytogenes* intracellular growth

RAW264. 7 cells were seeded at a density of 6 × 10^6^ cells per 24-well tissue-culture plate and allowed to adhere overnight. iBMDM were seeded at a density of 4 × 10^6^ cells per 24-well plate, allowed to adhere for 4 hours and then activated overnight with 100 ng/ml of LPS and interferon-γ. Where indicated, cells were pre-treated with 10 mM NAC, 500 µM L-glutathione reduced (GSH), 10 µM GSK-PERK inhibitor or 50 µM 4µ8c Ire1 inhibitor for 30 minutes followed by a 30 minute treatment with 3. 5 µM of DUB^inh^ or equivalent volume of DMSO (vehicle control). DUB^inh^ were removed from cells by changing the medium. Macrophages were then infected with *L. monocytogenes* at a multiplicity of infection (MOI) of 1 for 30 minutes. The inoculum was removed by washing cells with PBS three times, and cells were incubated in medium containing 10 µg/ml of gentamicin. Cells were collected at 6h post-infection in 1 ml of 0. 1% Triton X-100, serially diluted and plated on Luria-Bertani (LB) agar for enumeration. Colony forming unit (CFU) were counted using the Acolyte plate reader and software (Microbiology International). Results were normalized to DMSO-treated cells.

### Virus infection and plaque assay

For all MNV infections, RAW264. 7 cells were seeded in 12-well plates at 5 × 10^5^ cells/well and grown overnight. Cells were incubated for 30 minutes at 37^°^C with 1 mL medium containing no reagent, 2. 5 μM C6, or equivalent volume of DMSO as vehicle control. Then cells were infected with MNV-1 (CW3 at pass 6) at an MOI of 5 and rocked on ice for 1 hour for viral adherence. Cells were then washed 3 times with ice cold PBS and warm medium was returned to the cells with no reagents, 2. 5 μM C6, or equivalent volume of DMSO. For NAC-and L-glutathione-treated samples, NAC was added to a final concentration of 10 mM and Glutathione at a final concentration of 500 μM in addition to C6 or DMSO after viral adherence on ice. Cells were then incubated at 37^°^C for 8 hours. Plates were frozen at −80^°^C overnight, then freeze-thawed twice. Plaque forming units measured by plaque assay as described before (77). For EerI experiments, cells were infected exactly as with C6 above, but at a concentration of 5. 0 μM both before and after MNV infection.

### RNAi knockdown

SiGENOME Smart Pool small interfering RNA (siRNA) for mouse p97/vcp and the non-targeting siRNA were purchased from Dharmacon, and the manufacturer’s protocol was followed for the delivery of siRNA into the cells. Briefly, RAW264. 7 cells were seeded in 24-well plates at 1. 5 × 10^6^ cells/well and grown overnight. Transient knockdown was done by transfecting the cells using the DharmaFECT reagent 4 (Dharmacon) and a final concentration of 50 nM siRNA per well. The medium was replaced 6h post-transfection. The cell viability was assessed at different time points after transfection by diluting 1 in 10 the WST-1 reagent (Roche) in the media. Cells were incubated for 1. 5h before reading the optical density of the supernatant at 440 and 600 nm. Results were normalized to NT siRNA-transfected cells.

### ROS and calcium measurements

Macrophages were seeded in 6-well plates and allowed to adhere overnight. Where indicated, cells were pre-treated with 10 mM NAC, 500 µM L-glutathione reduced (GSH) or 300 µM TUDCA for 30 minutes, followed by a 30 minute treatment with 3. 5 µM of DUB^inh^, 5 µM DUB^inh^ C6 or equivalent volume of DMSO (vehicle control). Alternatively, cells were treated for 1h with 10 µM EerI, or 2h with 5 µM MG132 or 0. 5µM epoxomicin. Culture medium was removed, cells washed three times with PBS and then incubated for 30 minutes with 5 µM CM-H_2_DCFDA (general ROS indicator) or 1 µM Fluo-4 AM (intracellular calcium indicator) in HBSS buffer containing 20 mM HEPES. After three PBS washes, cells were incubated for 30 minutes with fresh media. Cells were washed three times with PBS and removed from plates using cold PBS. For time course experiments, cells were treated with 3. 5 µM DUB^inh^ for 0. 5h or 10 µM EerI for 1h, washed and then incubated in fresh medium for the indicated time before staining. Peritoneal macrophages and splenocytes were kept in suspension and stained with 1. 25 µM or 2 µM of CM-H_2_DCFDA, respectively. Fluorescence was read on Fortessa or Canto flow cytometers (BD Biosciences) and data were further analyzed using FlowJo software. For all experiments using EerI, the intrinsic fluorescence of EerI was removed for calculation of the mean fluorescence intensity.

### *Xbp1* splicing assay

RNA was extracted using the RNeasy Mini Kit (QIAGEN) from samples treated with 10 µg/ml of tunicamycin for 4h or with 3. 5µM DUB^inh^ or equivalent volume of DMSO for the indicated time. cDNA synthesis was performed using 1 µg of RNA and *Xbp1* transcripts were amplified using following primers: forward 5´-GAACCAGGAGTTAAGAACACG-3´ and reverse 5´-AGGCAACAGTGTCAGAGTCC-3´. Amplification was done using using 30 cycles at 94^°^C for 1 min, 60^°^C for 1 min and 72^°^C for 1 min. PCR products were further digested with the restriction enzyme PstI and loaded onto a 3% agarose gel to differentiate between the spliced and unspliced forms of *Xbp1*.

### Statistical analysis

Results represent the mean and the corresponding standard deviation of the mean for at least three independent experiments (unless specified) as indicated in figure legends. Statistical analysis was performed using GraphPrism 7 software and unpaired, two-tailed student’s t test or one-way analysis of variance (ANOVA) with Dunnett or Bonferroni post-tests were used, as indicated in each legend. Our data mostly meet the assumption of the tests. No statistical methods were used to estimate sample size. Rather, sample size was chosen based on similar studies performed in the innate immunity field. No statistical methods were used to include or exclude samples.

## Data availability

All data presented in this article and the corresponding supplementary information file are available upon request from the corresponding author.

## Acknowledgments

We thank members of the O’Riordan and Wobus laboratories for suggestions. This work was supported by NIH/NIAID R21/R33 AI102106 to M. X. D. R. and C. E. W. M-E. C. was supported by fellowships from the Fond de recherche santé Québec (Fellowships **#**27881 and #31426).

## Author contributions

M-E. C., K. P and S. H performed experiments. M-E. C, K. P, C. E. W. and M. X. D. R. analyzed results.

H. D. S. provided reagents. M. E. C. and M. X. D. R. wrote the manuscript.

